# Splicing and epigenetic factors jointly regulate epidermal differentiation

**DOI:** 10.1101/325332

**Authors:** Sabine E.J. Tanis, Pascal W.T.C. Jansen, Huiqing Zhou, Michiel Vermeulen, Klaas W. Mulder

## Abstract

Epidermal homeostasis requires a continuous balance between progenitor cell proliferation and loss of differentiated cells from its surface. During this process cells undergo major changes in their transcriptional programs to accommodate new cellular functions. We found that transcriptional and post-transcriptional mechanisms underlying these changes are functionally connected and jointly control genes involved in cell adhesion, a key process in epidermal maintenance. Using siRNA-based perturbation screens, we identified novel DNA/RNA binding regulators of epidermal differentiation. Computational modeling and experimental validation identified functional interactions between the matrin-type 2 zinc-finger protein ZMAT2 and the epigenetic modifiers ING5, SMARCA5, BRD1, UHRF1, BPTF, SMARCC2. ZMAT2 is required to keep cells in an undifferentiated, proliferative state and quantitative proteomics identified ZMAT2 as an interactor of the pre-spliceosome. RNA-Immunoprecipitation and transcriptome-wide RNA splicing analysis showed that ZMAT2 associates with and regulates transcripts involved in cell adhesion in conjuction with ING5. Thus, joint control by post-transcriptional and epigenetic mechanisms is important to maintain epidermal cells in an undifferentiated state.

**Highlights:**

- Gene-perturbation screens identify a role for ZMAT2 in the control of human epidermal differentiation.
- ZMAT2 functionally interacts with known epigenetic regulators of epidermal differentiation.
- ZMAT2 interacts with the pre-spliceosome and transcripts involved in cell adhesion.
- ZMAT2 mediated splicing and epigenetic control jointly target an adhesion related transcriptional program in human epidermal stem cells.

## Introduction

As our understanding of gene expression regulation improves so does our awareness of its complex dynamics and timing. The transcription machinery and it’s co-factors, chromatin state, RNA splicing, miRNAs are only some of the myriad of processes contributing to the versatile functions of a cell. Regulation of gene expression is of particular importance for governing the delicate balance between proliferation and differentiation in stratified epithelial tissues such as the skin, breast or colon tissues. The high renewal rate in these tissues requires tight regulation of gene expression programs to avoid the generation of aberrantly behaving cells giving rise to diseases. Here, we use the skin as a model to study the contribution of genes to the regulation of gene expression programs governing epidermal proliferation and differentiation. The epidermal layer of the skin is completely replenished each month in a process driven by epidermal stem cells or keratinocytes, which reside attached on the basal membrane. Upon initiation of differentiation these cells stop proliferating, release their integrin anchors and move up through the different layers of the skin traveling upwards through the spinous and the granular layers. (Moreno-Layseca and Streuli, 2014). Eventually, they end up de-nucleated and heavily interconnected in the cornified layer, where they are shed off of the skin. This process is marked by, amongst others, downregulation of integrins and upregulation of differentiation genes such as envoplakin (ENV), periplakin (PPL) (Ruhrberg *et al.*, 1997), involucrin (INV) and transglutaminase (TGM1) (Eckert *et al.*, 2005). Throughout this transition they constantly change and fine-tune their expression programs using transcriptional and post-transcriptional processes in a differentiation state dependent manner.

These transitions in gene expression programs are controlled by regulatory mechanisms provided by epigenetic factors, transcription factors and post-transcriptional processes such as splicing. Epigenetic factors regulate chromatin accessibility through remodelling (BPTF, SMARCA5) or adding or removing histone or DNA modifications (ING5, UHRF1) (Mulder *et al.*, 2012). Open chromatin structures allow for transcription factors to bind their motifs and enable activation of gene expression programs. There are several transcription factors which have a known role in keratinocyte biology (AP1, ETS-family (Eckert *et al.*, 1997; P. Nagarajan *et al.*, 2010) to regulate the expression of differentiation markers (e.g. IRF6 (Biggs *et al.*, 2012)(Botti *et al.*, 2011a) and MAFB (Miyai *et al.*, 2016)). The emerging RNA transcript is then found by RNA-binding proteins that edit and protect it on its way to being translated into a protein. One of the most important RNA editing processes is RNA splicing, where introns are excised from the immature transcript generating a mature RNA transcript. More extensive editing via alternative splicing may be important in the skin, as there are several genes that display isoform specific expression in different keratinocyte cell states. For instance, dermokine, a gene that is highly expressed in the granular layer of the epidermis is heavily spliced, generating multiple isoforms with different functions (Naso *et al.*, 2007). Another example is desmoplakin (DSP) which has two major isoforms which seem to have distinct functions in controlling desmosomal adhesion in the skin (Cabral *et al.*, 2012).. These examples illustrate the importance of proper regulation of both transcriptional and post-transcriptional processes. However, our current understanding of how these processes coordinately govern epidermal biology is limited.

Here, we investigated the role of 145 putative DNA/RNA binding factors in human epidermal differentiation using siRNA-based perturbation screens. We identified the matrin type-2 zinc-finger protein ZMAT2 to be important to maintain cells in an undifferentiated state and as a splicing regulator of adhesion related transcripts. Moreover, computational predictions and experimental validation uncovered a previously unappreciated connection between epigenetic and post-transcriptional control of keratinocyte differentiation.

## Materials and methods

### Cell culture

Primary human keratinocytes were cultured on a mitotically inactivated J2-3T3 layer as described previously (Mulder *et al.*, 2012). Prior and during experiments where differentiation was induced, the cells were grown on keratinocyte serum-free medium (KSFM, Gibco) supplemented with 0.2 ng/mL epidermal growth factor (EGF, Gibco) and 30 ug/mL bovine pituitary extract (BPE, Gibco). Differentiation was induced using 10 μM AG1478 (Calbiochem), 200 ng/mL human recombinant BMP 2/7 (R&D systems) a combination of these or 10% fetal bovine serum (MP Bio) was added to the KSFM for 48 hours.

For colony formation assays, J2-3T3 cells were seeded at 100.000 cells per well in a 6 well format and inactivated using mitomycin C (2 μg/mL, Santa Cruz Biotechnology) as described previously (Mulder *et al.*, 2012). Transfected keratinocytes (500 cells) were seeded on top and grown for 2 weeks before fixation and immunostaining.

### siRNA nucleofection

Passage 3 lip or foreskin keratinocytes were grown to 70% confluency in KSFM before they were used for nucleofection using the Amaxa system (Lonza). The cells were collected and resuspended in cell line buffer SF at 2*10^5 cells per 18 uL, mixed with 2 uM siRNA. After transfection using program FF113, the cells were incubated for 10 minutes at room temperature. Finally, the transfected cells were resuspended in pre-warmed KSFM and dispensed over the culture plates manually.

### siRNA library

145 genes were selected based on differential expression and RPKM over 5 in a dataset published in Kouwenhoven *et al.*, 2015. Additionally, literature was manually searched to select those factors that had DNA binding capacities. A custom silencer select siRNA library was obtained from Ambion including 3 siRNAs per gene divided over 2 plates. The siRNA screens were performed using the pooled siRNA’s per gene.

### siRNA screening and data processing

Passage 3 foreskin keratinocytes were grown up to 70% confluency in KFSM for nucleofection. In addition to the custom library, siRNA controls were transfected; scramble siRNAs and siRNAs targeting transglutaminase I. After transfection, the cells were manually dispensed onto 20 96-well plates, generating quadruplicate plates for each differentiation inducing treatment plus control. The medium was refreshed the next day and after 72 hours and differentiation was induced (vehicle, AG1478, BMP 2/7, AG1478 + BMP 2/7 and serum) for 48 hours. The cells were fixed using 4% paraformaldehyde in PBS (10 minutes, at RT) and subsequently permeabilised using 0.1% Triton X-100 in PBS (10 minutes, at RT). The cells were blocked using 10% serum in PBS for 30 minutes at room temperature and subsequently stained with primary antibody targeting TGM1 (mouse anti-BC-1, 1:2000) for 1 hour at room temperature. Following 3 washes the cells were stained with mouse secondary antibody IR800 (IRDye 800CW, 1:2000) and DNA staining agent DRAQ5 (Biostatus, 1:4000) in blocking buffer for 1 hour at room temperature. After three washes a final volume of 100 uL PBS was added and the plates were scanned using the Licor Odyssey CLx system. The same settings were used for each screen.

Signals were quantified using the Licor Odyssey CLx software and used for subsequent analysis. The signal was normalized using the DRAQ5 DNA stain and the background signal measured in the TGM1 KD samples was subtracted. The data was further processed to z-scores based on knockdown readouts only, transforming the data to a standardized format enabling compilation of all the experimental data.

### Bayesian mixture model and network visualisation

The phenotypic profiles extracted from the screen, describing the effect of knockdown of a certain gene on TGM1 expression in 5 different conditions, are compared in this model. The network is visualized using the R-package RedeR (Castro *et al.*, 2012). The Bayesian mixture model is described in Wang *et al.* (2012). In short, we extracted the phenotypic profiles per gene from the z-score matrix, for which the cosine correlation between each gene pair was calculated. Under the assumption that there were three possible modes of functional association; positive, negative or a lack of association, there are three beta distributions in this model. To estimate the parameters α and β for the positive and negative distribution, maximum a posteriori probability (MAP) inference was applied with an uninformative prior (uniform Dirichlet priors) using an expectation maximization (EM) algorithm. For the parameters of the lack of association distribution, the data set is permutated 100 times to create a random dataset. The data is fitted to these distributions, generating a matrix with probabilities for each gene pair belonging to one of the three distributions. This matrix is subjected to network inference, looking at functional interactions and their signal to noise ratio (SNR: probability positive or negative association/probability lack of association). We only considered functional interactions that have a SNR of 10 or higher and checked the significance of the inferred connections with multiscale bootstrap resampling. Using the approximately unbiased method (sample size between 0.5 to 1.4), we avoided a bootstrap resampling bias. The displayed p-values are defined as 1-AU.

### Identification genetic interactions

Cells were transfected with 2 independent siRNAs per gene and the different combinations of these siRNAs in triplicate. RNA was isolated and CEL-seq 2.0 libraries were prepared. After sequencing and subsequent mapping using STAR the data was analysed using DESeq 2.0 to call differentially expressed genes versus siControl. We included all genes that were detected in at least 90% of the samples and significantly differentially expressed with p-value<0.05 in at least one comparison. Furthermore, samples in which more than 10% of these genes was not detected were excluded from further analysis to increase the robustness of the statistical analyses. For the remaining samples and genes, we calculated the fold change in expression compared to siControl (observed FC). The expected fold changes (NULL hypothesis in the absence of a functional interaction) were calculated using the product rule (Mani *et al.*, 2008) multiplying the FC for the individual knockdown samples for ZMAT2 versus the other genes. To estimate the variation of the NULL hypothesis, we performed this calculation for all combinations of independent siRNAs and replicates. The expected FC was then compared to the observed FC using a multiple testing corrected t-test (Benjamini-Hochberg, FCR <0.01). Interactions were labelled alleviating when the expected FC was higher than the observed FC and aggravating when the expected FC was lower than the observed FC.

### RNA extraction, RT-qPCR and expression profiling

RNA was isolated using the Quick-RNA^™^ MicroPrep kit from Zymo Research and subsequently 1 μg RNA was converted to complementary DNA. Thermo Maxima Reverse Transcriptase was used for reverse transcription and the resulting cDNA was diluted to ~5pmol/μL for qPCR using SYBR Green Master mix. Data was normalised using 18s signal as a control, which was included on each plate.

### Colony formation assay analysis

The cells were blocked using 10% serum in PBS for 30 minutes at room temperature and subsequently stained DNA staining DRAQ5 (Biostatus, 1:4000) in blocking buffer for 1 hour at room temperature. After three washes a final volume of PBS was added and the plates were scanned using the Licor Odyssey CLx system. The images were processed as described in (Buggenum *et al.*, 2017)

### IP-mass spectrometry

Passage 3 foreskin keratinocytes were grown up to 70% confluency in KFSM before harvesting and counting. Nuclear extracts were prepared by adding 5 volumes of buffer A (10 mM HEPES KOH pH 7.9, 1.5 mM MgCl_2_, 10 mM KCl) and incubation of 1 hour on ice. The cells were centrifuged for 5 minutes at 450g before lysis by dounce homogenization in 2 volumes cell pellet buffer A with 0,5% NP40 and protease inhibitors. Cytoplasmic and nuclear extract were separated by centrifugation at 3200g for 15 minutes and collection of cytoplasmic fraction as supernatant. The nuclear pellet was washed with 10 volume PBS and centrifuged at 3200g for 5 minutes. Nuclei were resuspended in 2 volumes extraction buffer C (420 mM NaCl, 20 mM HEPES KOH pH 7.9, 20% v/v glycerol, 2 mM MgCl_2_, 0.2 mM EDTA, 0.5% NP40, 0.5 mM DTT, protease inhibitors) and incubated 2 hours rotating at 4°C. The extract was centrifuged for 30 minutes at 14.000 rpm and the nuclear extract was collected. Protein concentration was determined using a Bradford assay.

500 ug nuclear extract was used in an overnight immunoprecipitation with 3 μg α-ZMAT2 (DSHB, PRCP-ZMAT2-1E5) in a total volume of 500 μL. This was mixed with 25 uL Dynabeads magnetic beads (Invitrogen) and incubated for 4 hours rotating at 4°C. After incubation, beads where washed three times using buffer A, two times using PBS, once using ABC buffer (25 mM ammonium bicarbonate) before final resuspension in 50 μL ABC buffer. Overnight digestion was performed using 350 ng trypsin where after samples were desalted using C18 stagetips (Rappsilber, Mann and Ishihama, 2007).

The tryptic peptides were separated in a 120 minute acetonitrile gradient (7% to 32%, step-wise increase up to 95%) on an Easy nLC 1000 (Thermo Scientific) connected online to a Orbitrap Fusion Tribrid mass spectrometer. MS and MS/MS spectra were recorded as described in van Buggenum et al 2018. Data analysis was performed as described before (Smits *et al.*, 2013) using MaxQuant version 1.5.1.0 (Cox and Mann, 2008) and protein database UniProt_201512\HUMAN. Perseus was used to filter the data and the plots were made in R.

### IP-RNA sample preparation

Passage 3 foreskin keratinocytes were grown up to 70% confluency in KFSM before harvesting and counting. After preparing the nuclear extract and determining the protein concentration by Bradford assay,the extract was divided over samples for input RNA isolation and immunoprecipitation in triplicate. For input RNA, RNA was isolated directly from the nuclear extract and stored at −20°C. Immunoprecipitation was performed as described in the IP-MS paragraph up to washing the beads. RNA lysis buffer was added to the beads and incubated for 5 minutes before removing the beads from the sample using the magnetic tray. RNA was isolated from the supernatant as indicated in the protocol provided with the Quick-RNA^™^ MicroPrep kit (Zymo Research). The isolated RNA was further processed for sequencing using the CEL-seq 2.0 library prepation protocol and subsequent analysis.

### Epifluorescence localisation ZMAT2

Cells were transfected with scramble siRNAs or siRNAs targeting ZMAT2 seeded in 96-well glass bottom plates and cultured for 72 hours as indicated. Cells were washed, fixed using 4% paraformaldehyde for 10 minutes and subsequently permeabilised using 0.3% triton in PBS for 10 minutes. The cells were pre-blocked using 0.2M glycine for 20 minutes and blocked for 1 hour using 1% BSA. Cells were incubated overnight at 4°C with primary stain, which included a no-primary-antibody control and 4 μg/mL α-ZMAT2. After three wash steps with PBS cells were stained with secondary antibody Alexa 488 rabbit α-mouse (1:2000) for 1 hour and DAPI (1:500) in the last 10 minutes. Cells were washed 3 times and imaged on a Leica DMI6000B automated high-content microscope.

### Library preparation RNA sequencing

RNA sequencing libraries were prepared as described in the protocol supplied with the Illumina KAPA RNA HyperPrep Kit with RiboErase (HMR) – (KR1351), starting from 500 ng. The concentration of the libraries was determined using the Denovix HS dsDNA assay and library quality was checked using the Bioanalyser platform (Agilent). Samples were sequenced using the NextSeq500 (Illumina) platform.

### CEL-seq 2.0

CEL-seq 2.0 libraries were generated using the protocol from (Hashimshony *et al.*, 2016) with the following adaptations. 100pg purified RNA was directly added to a reverse transcription mixtures containing Maxima H Minus (ThermoFisher) reverse transcriptase and CEL-seq 2.0 primers with a 6-nucleotide sample barcode and 8-nucleodide UMI. After reverse transcription samples were pooled and purified using AmpureXP beads (Beckman Coulter). Second strand synthesis was performed according to the original protocol, libraries were amplified using 2 consecutive PCR reactions.

### Data analysis sequencing data

CEL-seq 2.0 data was processed using the Yanai pipeline (Hashimshony *et al.*, 2016) up to count tables which were further processed using DESeq 2.0. RNA sequencing data from KAPA RNA HyperPrep kit generated libraries was aligned (hg38, annotation: ensembl genecode basic annotation V27) and sorted using STAR and analysed using DEXSeq. For the analysis using MISO the sequencing data was aligned to hg19 using STAR whereafter the collection annotations under the Human genome (hg19) alternative events v1.0 was used to call alternative events. Data is deposited in GEO under accession code GSE114529

### Expression ZMAT2 in different tissues

Expression of ZMAT2 and other spliceosome components was taken from the EMBL expression atlas, which extracts its raw data from the PRIDE proteomics database. Website open source database: https://www.ebi.ac.uk/gxa/home/

## Results

### RNAi screens identify nucleic acid binding proteins involved in human keratinocyte renewal and differentiation

To investigate the role of putative nucleic acid binding factors in the regulation of epidermal stem cell renewal and differentiation, we performed siRNA-based knock-down screens (Figure 1A). For this, we selected 145 genes based on their expression levels in keratinocytes (RPKM>5), differential expression during differentiation (Supplemental Dataset 1, (Kouwenhoven *et al.*, 2015)) and DNA or RNA binding potential. To test the contribution of these genes to the process of differentiation, we silenced each gene using a pool of three independent siRNAs in triplicate and subsequently induced differentiation. Transfected cells were cultured in different conditions for 48 hours to induce differentiation with the EGF inhibitor AG1478, BMP 2/7, a combination of these compounds, 10% fetal bovine serum and a vehicle control, as previously described (Mulder *et al.*, 2012). These treatments lead to robust induction of the terminal differentiation marker transglutaminase I (TGM1) (Mulder *et al.*, 2012). Endogenous TGM1 protein was quantified using a fluorescent In-Cell-Western assay with an antibody (BC.1), whose specificity was confirmed using two independent siRNAs (Figure S1A). To account for differences in cell number among the knock-down populations in the screen, TGM1 measurements were normalised on DNA content (DRAQ5 signal) for each well. After z-score transformation, the results were compiled into a single dataset covering all 145 genes, 5 conditions and replicates (supplementary dataset 1). High correlation between replicates highlighted the reproducibility of our findings and the quality of our dataset (Figure S1B). Using a random selection of 11 genes, we estimate the median knock-down efficiency to be 88% (range: 28-93%) and the false negative rate at below 10% (Figure S1C). In addition, we performed deconvolution experiments where the individual siRNAs were tested in parallel with the pool of 3 siRNAs that was used in the screen. This indicated that 71-87% of the siRNA pools contained at least 2 siRNAs that recapitulated screen results (Figure S1D), arguing against widespread off-target effects. Taken together, our dataset constitutes a quality resource of nucleic acid binding factors to further characterise for their role in epidermal self-renewal and differentiation.

**Figure 1:**
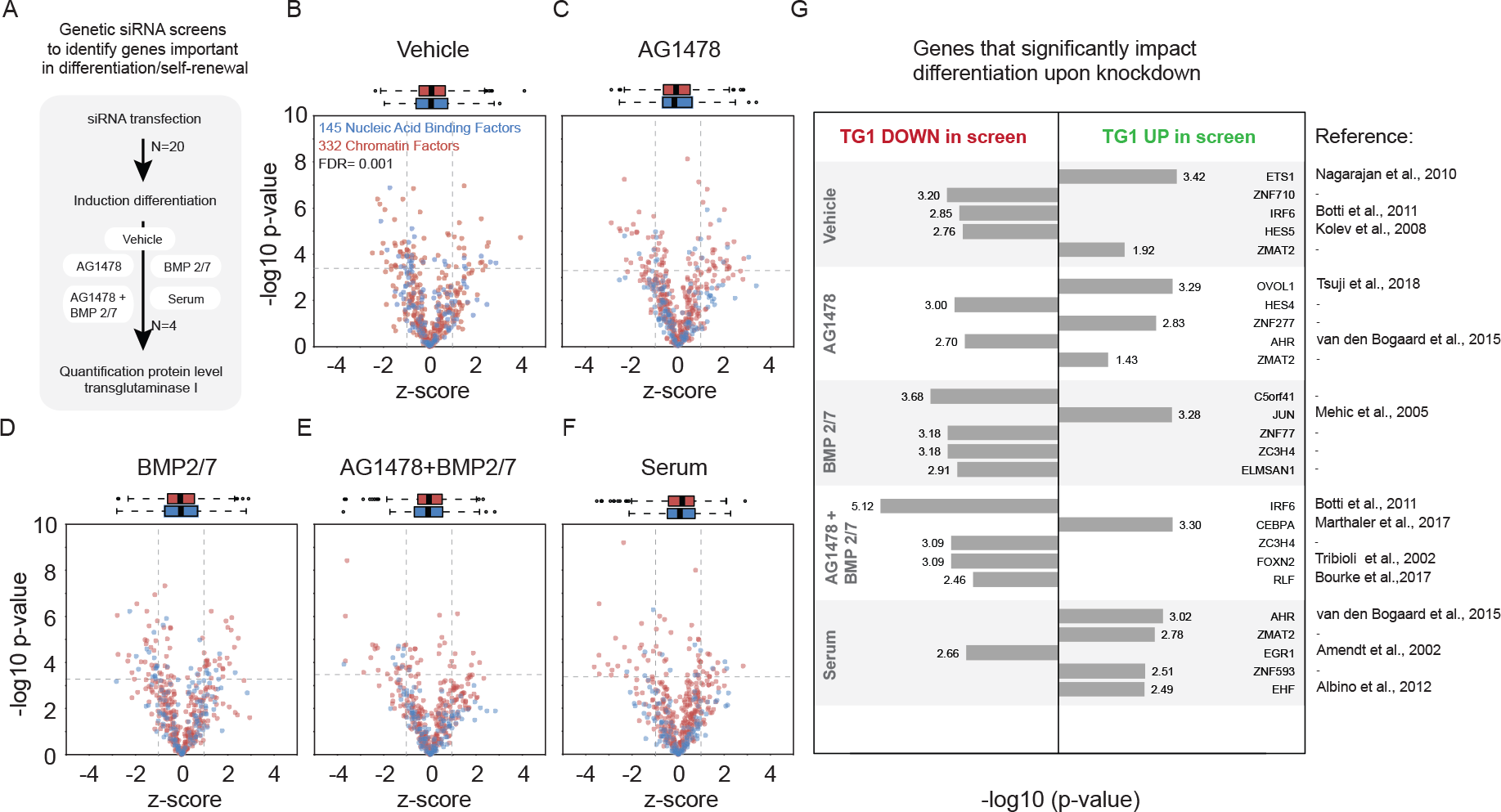
siRNA screens reveal novel nucleic acid binding factors involved in epidermal differentiation. (A) Schematic set-up of the screen; siRNA transfection (N=20) is followed by induction of differentiation (N=4, 5 conditions, 48h) and quantification of protein levels of transglutaminase I. (B) – (F) Volcanoplots describing the effects of siRNA based perturbation on TGM1 protein expression (x-axis) and their significance (y-axis) (N=3). Data from chromatin factors (CF) is displayed in red, nucleic acid (NA) binding factors in blue. Bars on top of graphs indicate spread of z-scores for each condition. FDR=0.001 (Benjamini-Hochberg) (G) Top 5 most significant genes in each condition, of which knockdown results in down or up regulation of transglutaminase I levels in the 5 different conditions. Significance is added as −log_10_(p-value). Literature references for genes with a known function in keratinocyte biology are highlighted under Refs (N=3).

From this data we identified putative nucleic acid binding factors that significantly affected TGM1 levels (Benjamini Hochberg (BH) FDR < 0.001) compared to the average across all siRNAs in the screen in any of the conditions (Supplemental Dataset 1, Figures 1B-F, blue datapoints). The normalised effect size per gene is plotted on the x-axis, whereas the statistical significance is depicted on the y-axis. These representations of the data highlight the factors that modulate differentiation in the top left and top right quadrants, respectively. Notably, the effects and significance of individual factors were condition dependent, indicating that not all identified factors play equivalent roles in the different conditions tested (Supplemental Dataset 1, Figure S2). This is represented by the partial correlation between the effect sizes across the conditions (Figure S2, scatterplots in bottom half) and the significant differences when comparing different conditions (Figure S2, vulcanoplots in top half). In total, our screens revealed 57 genes that display a significant effect in at least one of the conditions (supplemental dataset 1) indicating that our experiment identified factors that have a potential regulatory role in keratinocyte differentiation.

To explore these individual hits, we first selected the top 5 most significant genes (BH FDR <0.001) for each condition (Figure 1G). These represent 21 distinct nucleic acids binding factors, one third of which (7) have previously been implicated in epidermal renewal/differentiation, confirming the validity of our approach. We sought to experimentally verify that the TGM1 measurements in our screen indeed represented bona fide differentiation and not solely regulation of TGM1 levels. To this end, we selected IRF6 and ETS1 as exemplars and used RT-qPCR as an alternative read-out of the expression of differentiation and basal cell markers after siRNA mediated silencing. IRF6 has been shown to be important for the expression of genes important for epidermal differentiation (Botti *et al.*, 2011a). In addition, ETS1 is involved in keeping the cells in undifferentiated state by repressing genes involved in the formation of the cornified envelope (Priyadharsini Nagarajan *et al.*, 2010). RT-qPCR analysis confirmed the effects of the knock-down of these genes on TGM1 protein levels in our experiments (Figure S3). Moreover, we found that silencing IRF6 resulted in lower expression of the differentiation markers PPL, EVPL and IVL, whereas ETS1 silencing resulted in induction of these differentiation markers and a reduction in the basal cell markers ITGA6 and ITGB1. This is in line with literature where IRF6 is thought to regulate differentiation and ETS1 is important in the self-renewing state. These results also confirm that the effects of silencing IRF6 or ETS1 in our screen indeed reflect the cellular differentiation state and not merely misregulation of TGM1 expression.

### ZMAT2 functionally interacts with epigenetic regulators of epidermal cell renewal

A powerful feature of the approach we took is that our siRNA based screen set-up allows us to combine the current results on 145 DNA/RNA binding factors with published data characterising ~330 epigenetic regulators (Mulder *et al.*, 2012). Moreover, it enables us to use a previously developed a Bayesian statistical framework that reveals genes that functionally interact (Wang *et al.*, 2012). This means identifying sets of genes that share functionality through regulation of similar cell biological processes, in this case epidermal differentiation. Applying this statistical approach on a combined epigenetic and DNA/RNA binding factor dataset should therefore allow identification of novel functional interactions among these groups of genes, leading to insights into their joint regulation of epidermal biology. The data distribution and variation of the two datasets are highly comparable, allowing us to combine the two datasets for further analysis (Figure 1B-F boxplots, Figure S4). This resulted in a rich dataset comprising 473 genes describing their effects on the expression of TGM1 in 5 conditions. Application of the Bayesian network algorithm to this joint dataset revealed strong predicted functional interactions between ZMAT2, a matrin-type 2 like zinc-finger with potential DNA or RNA binding capacity, and components of a previously identified network of epigenetic regulators involved in epidermal renewal (Mulder *et al.*, 2012, Figure 2A, full network in Figure S5). This subnetwork contained multiple members of different protein complexes representing diverse epigenetic mechanisms such as MORF complex members ING5 and BRD1, NURF complex members BPTF and SMARCA5, as well as SMARCC2 and UHRF1 (Mulder *et al.*, 2012). This implies that ZMAT2 plays a role in epidermal differentiation in conjunction with these epigenetic modifiers.

**Figure 2:**
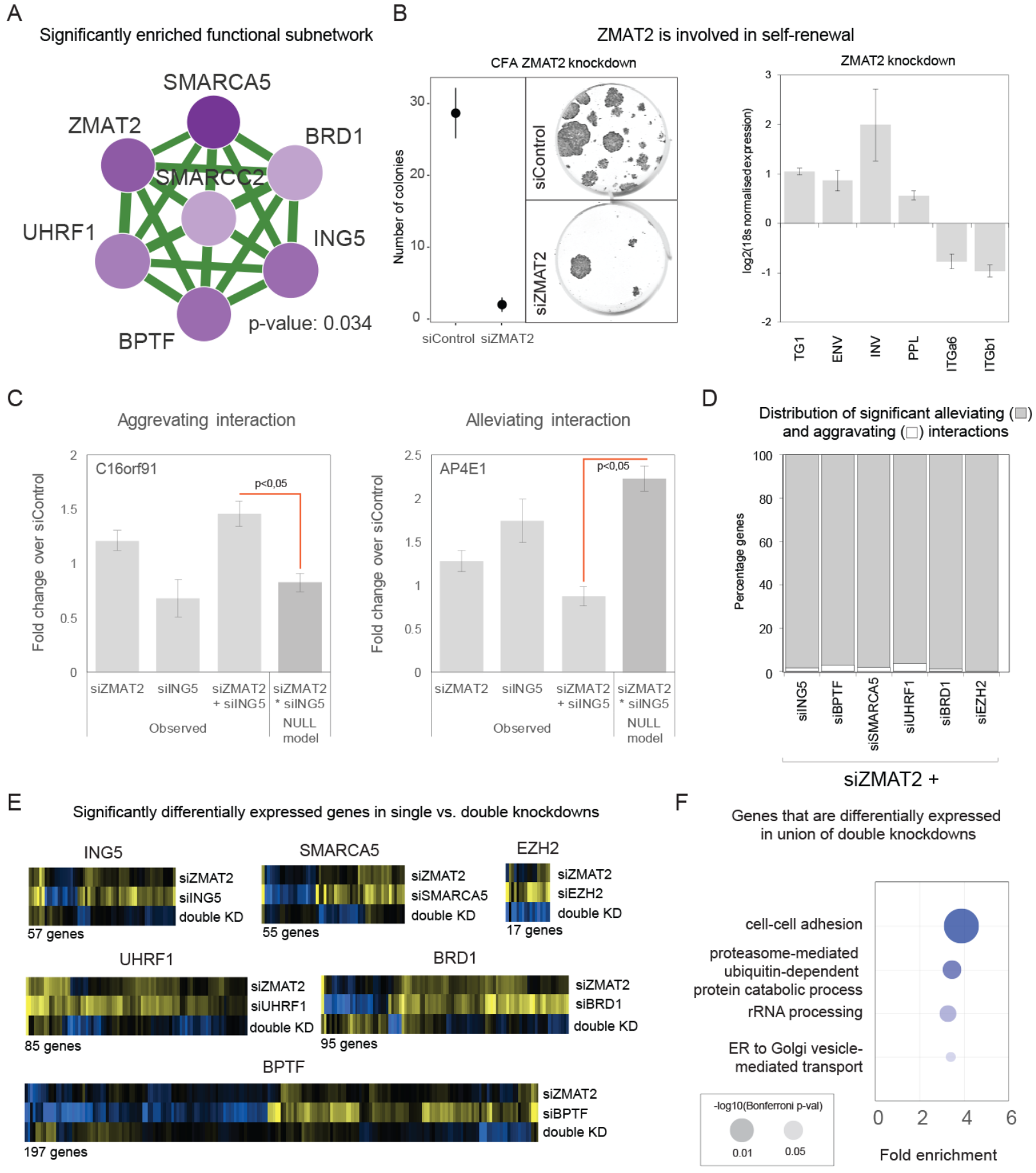
Bayesian network prediction reveals a functional interaction network between epigenetic factors and ZMAT2. (A) The significant subnetwork predicting functional interactions between epigenetic factors ING5, SMARCA5, SMARCC2, BRD1, BPTF and UHRF1 and zinc-finger ZMAT2. (N=3, an empirical p-value was determined using a bootstrapping approach) (B) Colony formation assays (N=3) and RT-qPCR based characterisation (N=3) of cells with perturbed ZMAT2 expression indicates a role in self-renewal. (C) Examples for an aggravating and alleviating functional interaction, taken from the total dataset. (siZMAT2, N=6; siING5, N=6; siZMAT2+siING5, N=12; NULL model, N=36). Errorbars indicate SEM. (D) Distribution of the significant aggravating and alleviating functional interactions for each of the combined knockdown experiments (E) Heatmaps visualising differential expression within the different samples (F) GO term analysis on the union of all significantly differentially expressed genes within the knockdown combinations (DAVID).

These predicted functional interactions prompted us to functionally characterise ZMAT2 further. RT-qPCR analysis showed that silencing ZMAT2 in keratinocytes resulted in increased expression of differentiation markers (PPL, EVPL, INV, TGM1) and concordant down regulation of basal-cell markers ITGB1 and ITGA6 (Figure 2B). In addition, depletion of ZMAT2 resulted in a strong reduction of the number and size of clones in colony formation assay (Figure 2B), reflecting a loss of cell renewal and proliferation capacity. These results indicated that ZMAT2 at least plays a role in maintaining epidermal cells in an undifferentiated, proliferative state and confirmed our primary screen results.

The Bayesian mixture model predicted that ZMAT2 functionally interacts with the epigenetic regulators ING5, SMARCA5, BPTF, UHRF1, BRD1. We aimed to confirm these predicted interactions using a double knock-down strategy. In this way, true functional (genetic) interactions can be identified (Mani *et al.*, 2008; Costanzo *et al.*, 2010; Horn *et al.*, 2011; Roguev *et al.*, 2013) by comparing the quantitative effects of combinatorial knockdowns on global gene expression with effects that can be expected based on the individual knock-down (Horn *et al.*, 2011). In cases where two genes are independent (i.e. do not functionally interact), the observed phenotype in the double knock-down is equivalent to the product of the individual knock-down phenotypes (Costanzo *et al.*, 2010; Horn *et al.*, 2011; Roguev *et al.*, 2013). Using this ’product-rule’ as the null hypothesis, genetic interactions are statistically defined as alleviating (where the observed phenotype of the combined knock-down is less pronounced than expected) or aggravating (where the observed phenotype is greater than expected), respectively (examples Figure 2C). Generally, alleviating interactions occur between genes involved in the same process/pathway, whereas aggravating interactions tend to be associated with functionally redundant processes (Costanzo *et al.*, 2010; Roguev *et al.*, 2013).

In previous work we confirmed that the epigenetic regulators within the predicted subnetwork (Figure 2A) display true functional interactions (Mulder *et al.*, 2012). Therefore, we decided to experimentally test the predicted interactions between ZMAT2 and ING5, SMARCA5, BPTF, UHRF1 or BRD1 using a combined knock-down approach. As a control we included EZH2, which is only peripherally associated with the epigenetic factors in this network (Mulder *et al.*, 2012). To obtain a detailed quantitative phenotype representing the cell state, we performed RNA expression profiling after silencing each of these genes individually, as well as in combination with ZMAT2. Two independent siRNAs targeting each gene were used in all possible permutations in triplicate. We analysed a total of 156 knockdown samples using a modified CEL-seq 2.0 method that enables high-throughput 3′-end tag-counting RNA-sequencing (see materials and methods for details). It is important to note that the relatively shallow RNA-sequencing we performed does not allow us to confidently interpret the molecular mechanisms underlying these interactions but is powerful for discovery of functional interactions based on RNA expression profiles. We examined the data using DESeq 2.0 and selected transcripts that were significantly differentially expressed (p<0.05) compared to control siRNA-transfected cells. After quality control and filtering (see materials and methods), we identified transcripts that displayed statistically significant functional interactions based on the product-rule criteria. For each of the factors we silenced in combination with ZMAT2, between 55 and ~200 genes displayed such functional interactions (Figure 2E). In contrast, the peripherally associated factor EZH2 showed only 17 functional interacting transcripts. Notably, the vast majority (>95% in all cases) of the functional interactions we identified were alleviating interactions, suggesting that ZMAT2, ING5, SMARCA5, BPTF, UHRF1 and BRD1 function in a similar process/pathway (Figures 2D and S6). GO-term enrichment analysis on the genes contributing to the observed functional interactions revealed an overrepresentation of factors involved in cell adhesion, a process required for anchoring epidermal stem cells to their niche (Figure 2F). These findings show that we were able to experimentally validate the predicted functional interactions between these epigenetic factors and ZMAT2 and that this group of genes jointly regulate an expression program linked to cell adhesion.

### ZMAT2 associates with the pre-spliceosome in epidermal keratinocytes

In order to gain insight into the potential molecular role of ZMAT2 in epidermal cell biology, we aimed to identify the proteins it associates with using immuno-precipitation and quantitative mass spectrometry (IP-MS, Figure 3A). Reminiscent of the epigenetic factors, the ZMAT2 protein localizes to the nucleus, as determined by immuno-fluorescence microscopy (Figure S7). Antibody specificity was confirmed using siRNA-mediated knockdown, resulting in decreased nuclear ZMAT2 staining (Figure S7). Next, we performed immuno-precipitation using control IgG (non-targeting/background binding) and antibodies targeting endogenous ZMAT2 from a nuclear extract. Following IP, sample prep and desalting, these samples were subjected to label-free quantitative mass-spectrometry to identify associated proteins. The data is plotted as enrichment over IgG control versus the significance over the three replicates with significant interactors located in the right upper quadrant (Figure 3B). Both Gene Ontology and protein interaction database analysis of the interacting proteins revealed that ZMAT2 interacts with components of the (pre-) spliceosome (Figures 3C,D). These findings are in line with crystal structures of the budding yeast pre-spliceosome, showing that the putative ZMAT2 ortholog snu23 associates with the B* (pre-catalytic) complex (Plaschka, Lin and Nagai, 2017; Ulrich and Wahl, 2017). Our biochemical experiments identifying ZMAT2 as an interactor of the pre-spliceosome in human keratinocytes, together with our screen and validation analysis implicate RNA splicing as a potential key regulatory process in epidermal renewal and differentiation.

**Figure 3:**
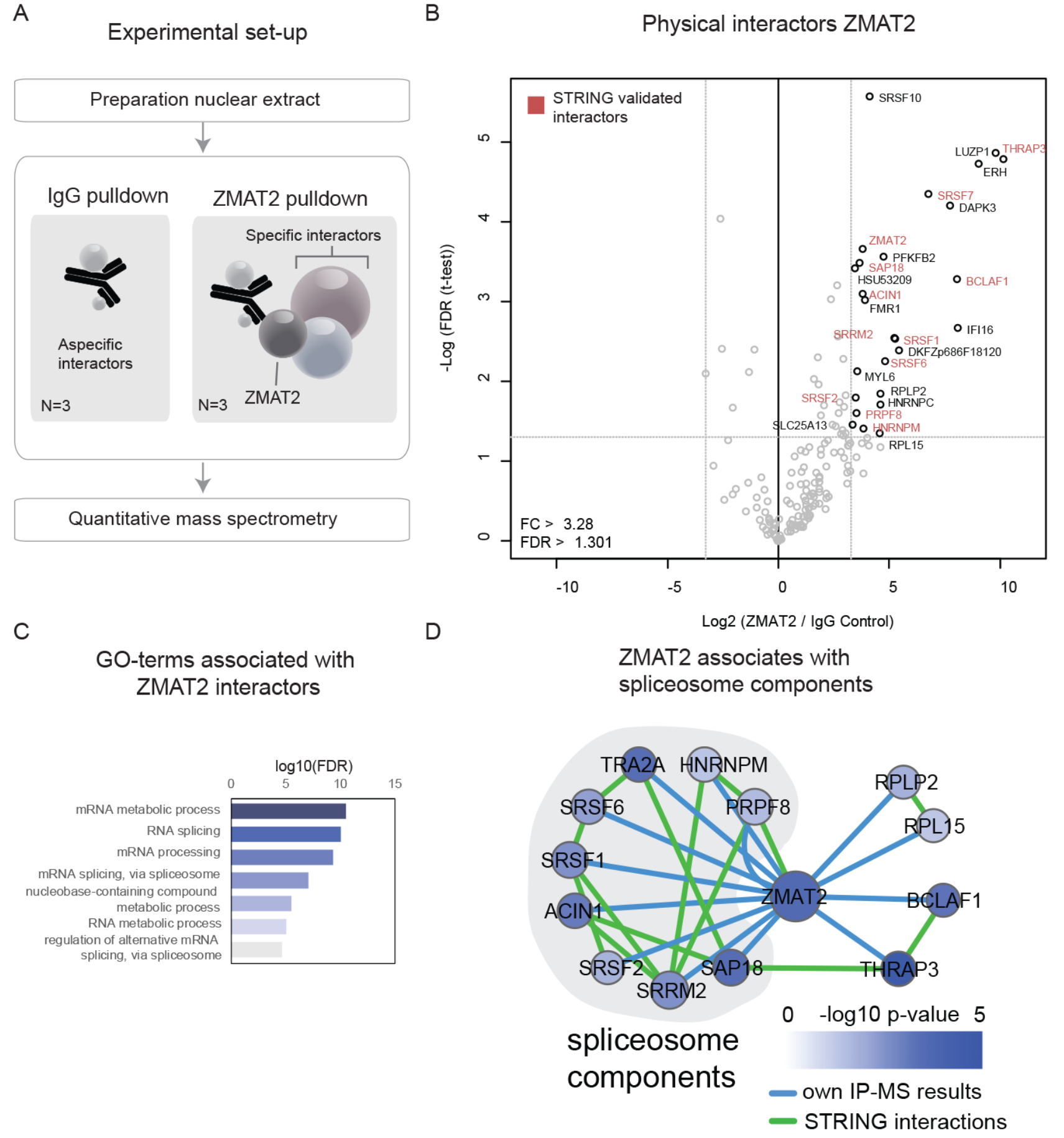
ZMAT2 interacts with components from the pre-spliceosome. (A) Schematic representation of the experimental set-up. Nuclear extract was prepared, one which IgG and ZMAT2 pulldowns were performed in triplicate. The peptides were quantified using label free mass-spectrometry. (B) Volcanoplot displaying the physical interactors of ZMAT2, with the log2 fold enrichment of ZMAT2 associated proteins over aspecific IgG bound proteins plotted against their significance (N=3). (C) GO term analysis (STRING) on proteins associated with ZMAT2. (D) STRING interactions versus experimentally found interactions with their significance displayed in blue.

### ZMAT2 associates with a subset of cellular transcripts involved in epidermal biology

Even though mRNA splicing is a ubiquitous process, we wondered if ZMAT2 containing spliceosomes display selectivity towards specific transcripts, or whether ZMAT2 is associated with essentially all transcripts present in the cell. An indication for a more selective role for ZMAT2 stems from the observation that the ZMAT2 protein is expressed in tissue specific manner, whereas other pre-spliceosome components show much more ubiquitous expression in the human body (Figure S8). Another indication comes from the GO-term analysis on the genes that are regulated by the functionally connected genes in our double knockdown experiments, where we found an enrichment of adhesion-related genes (Figure 2F). To identify ZMAT2-associated transcripts, we performed RNA-immunoprecipitation-sequencing (RIP-seq) with non-specific control IgGs and ZMAT2 specific antibodies from nuclear extracts. RNA-IPs were performed in triplicate and the isolated RNA was used to generate RNA-sequencing libraries using the modified CEL-Seq 2.0 protocol (Figure 4A). This specific form of library-prep is limited to the evaluation of poly-adenylated transcripts, but does allow analysis of the low amount of material retrieved after immunoprecipitation. To assess the specificity of enrichment, the recovered transcripts were compared to RNA-sequencing of the input material. Although the same amount of input material for IgG and ZMAT2 pull-downs was used, the IP with the ZMAT2 antibodies yielded substantially more RNA compared to the control IgGs, as can be expected from its association with the spliceosome (Figure 4B). The library complexity (number of different transcripts detected per 1000 reads) of the transcripts associated with the control IgGs reflected that of the input, reflecting non-specific interactions. In contrast, the complexity of the ZMAT2 associated RNAs was lower, suggesting that ZMAT2 associates with a subset of the total available transcripts, rather than binding RNAs indiscriminately (Figure 4C).

**Figure 4:**
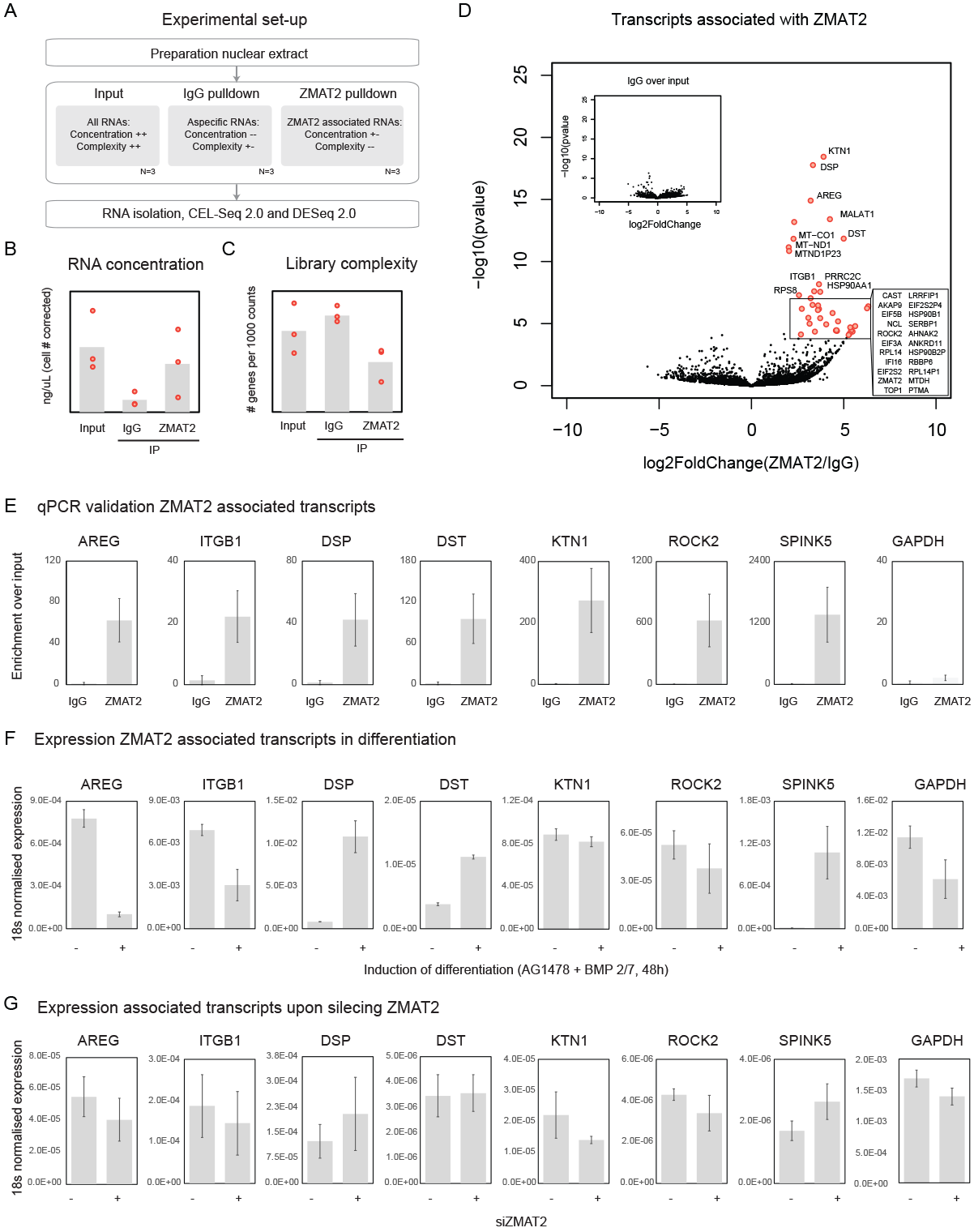
Adhesion related transcripts are associated with ZMAT2. (A) Schematic representation of the experimental set-up. Nuclear extract was prepared and normal immunoprecipitation procedure was performed, up to the wash steps. RNA was isolated directly from the beads and prepped for sequencing using CEL-Seq 2.0. (B) RNA concentration samples before and after the IP procedure. Corrected for cell numbers, in ng/μL (N=3). (C) Complexity of the CEL-Seq 2.0 libraries after sequencing. Number of identified genes per 1000 counts (N=3). (D) Volcanoplot displaying the transcripts that are significantly enriched in the ZMAT2-IP sample over the IgG-IP sample as called by DESeq 2.0. Inset represents significant hits identified in the IgG sample (N=3). (E) qPCR validation of ZMAT2 associated transcripts showing the enrichment over the input sample (N=3, error bars indicate S.E.). (F) Expression of the ZMAT2 associated transcripts is differential during induction of differentiation using EGF inhibitor AG1478 (10 μM) and BMP 2/7 for 48 hours (N=3, error bars indicate S.E.). (G) Expression of ZMAT2 associated transcripts upon silencing ZMAT2 (70% KD, 5 days of culture) as determined via qPCR (N=3, error bars indicate S.E.).

Next, we used statistical analysis (DESeq 2.0) to identify significantly enriched transcripts for both the IgG and the ZMAT2 samples. None of the RNAs identified in the control IgG IP were significantly enriched compared to input, indicating that these indeed represented non-specific interactions (Figure 4D, inset). In contrast, nearly 100 transcripts were significantly enriched in the ZMAT2 pull-down (Figure 4D). We choose to use the IgG data as a comparison for the ZMAT2 sample to ensure that we were comparing samples that had been processed in an identical fashion. Interestingly, many of the transcripts that were specifically and strongly associated with ZMAT2 are involved in cell-adhesion (e.g. ITGB1 (Levy *et al.*, 2000), DSP (Cabral *et al.*, 2012), DST (Michael *et al.*, 2014) and ROCK2 (Lock *et al.*, 2012) and proliferation processes (e.g. AREG (Stoll *et al.*, 2016)). Because both the functional interaction and RNA-IP experiments point towards regulation of adhesion we decided to focus on these transcripts. We selected 7 of these transcripts for validation by RNA-IP followed by RT-qPCR and included GAPDH as a negative control. This confirmed that all 7 transcripts are indeed highly enriched in the ZMAT2 pull-downs whereas GAPDH is not, confirming the selectivity of RNAs associated with ZMAT2 (Figure 4E). The lack of enrichment of GAPDH, one of the most abundantly expressed transcripts, in the ZMAT2 pull-down indicated that the identified associations are not solely based on transcript abundance. Moreover, of these 7 transcripts ITGB1, AREG, DSP, DST and SPINK5 are differentially expressed during differentiation (Figure 4F), suggesting that, at least some of, the ZMAT2 associated transcripts are also regulated at the transcriptional level. However, ZMAT2 itself does not seem to be involved in regulation of their expression, as silencing ZMAT2 does not affect the expression of ITGB1, ROCK2, AREG and DST. Differential expression of KTN1 (down regulated) and DSP and SPINK5 (up regulated) are likely an effect of ZMAT2 silencing on the differentiation state of the cell (Figure 4G and 2B). Together, these experiments indicate that ZMAT2 associates, presumably through its interaction with the pre-spliceosome, with a specific subset of transcripts in human epidermal cells.

### ZMAT2 silencing leads to differential exon usage of selected genes

To investigate whether ZMAT2 silencing indeed affects splicing in our cells, we performed full-length transcript RNA-sequencing on ZMAT2 knockdown samples in triplicate (Figure 5A). We included the ZMAT2 interacting pre-spliceosome component SRSF1 as a positive control for disruption of splicing (Figure 3B). SRSF1 is expected to lead to defects in splicing as it is involved in the first steps of assembly of the spliceosome by enhancing the binding of the U1 snRNP on the pre-mRNA (Kohtz *et al.*, 1994). DEX-Seq (Anders, Reyes and Huber, 2012) was used to calculate and statistically assess differential exon usage for all identified transcripts to detect altered splicing activity. This revealed that fewer genes are affected in their splicing by silencing of ZMAT2 compared to silencing SRSF1 (6.6% versus 25.9% of all identified transcripts, respectively. Figure 5B). Moreover, ZMAT2 regulated a smaller proportion of all potential exons in the transcriptome (0.18%), compared to SRSF1 (1.53%) (Figure 5C). Although it transpires that neither factor is essential for splicing per se, these findings indicate that ZMAT2 regulates a more restricted set of transcripts compared to SRSF1. Exon usage of about 200 transcripts was affected by both ZMAT2 and SRSF1. Of these 200 transcripts, the exons that were differentially regulated by these factors were predominantly mutually exclusive (Figure 5D), further supporting the notion of specific functions for ZMAT2 compared to the core pre-spliceosome component SRSF1 in epidermal keratinocytes. An exemplar of this is the DMKN gene which we experimentally validated using exon specific RT-qPCR (Figure S9). To investigate if there were differences in usage of specific splicing mechanisms between ZMAT2 and SRSF1, we used the MISO package, which enables a differential splicing analysis based on annotation files describing different splice events. This did not reveal major differences in splicing mechanisms (skipped exons (SE), intron retention (RI), mutually exclusive exons (MXE), alternative 5’ splice sites (A5SS), or alternative 3’ splice sites (A3SS)) affected by ZMAT2 compared to SRSF1 (Figure S10). Together, these experiments suggest that ZMAT2 regulates exon usage of a specific subset of transcripts, rather that functioning as a global splicing regulator in epidermal stem cells. We would like to note that our splicing analysis is unlikely to be fully comprehensive, as the there is some enrichment of abundant transcripts in our data, precluding interrogation of low expressed genes (Figure S11).

**Figure 5:**
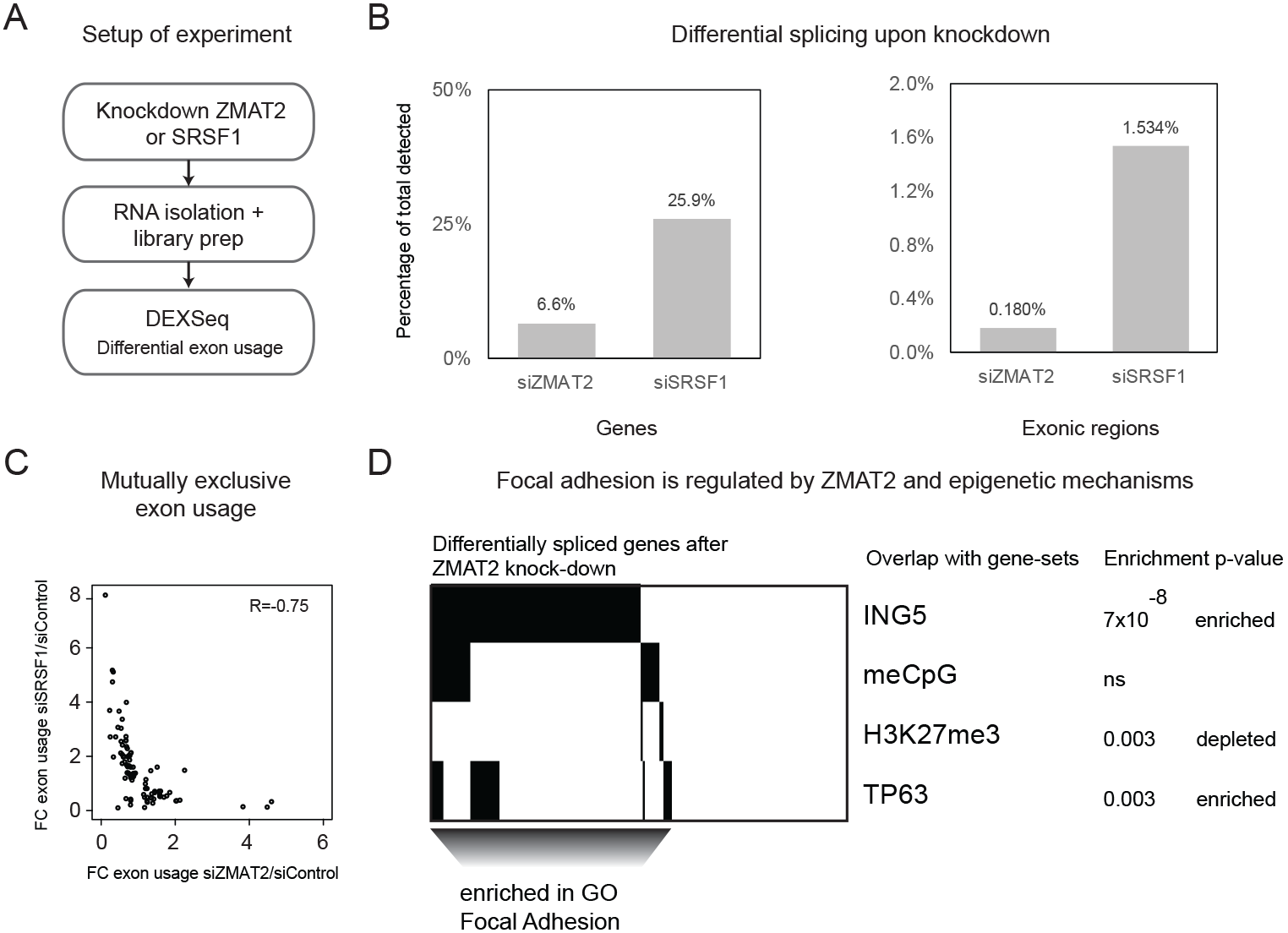
ZMAT2 regulates a similar gene set as the functionally interacting epigenetic factors through modulation of RNA splicing. (A) Schematic representation of the experimental set-up. ZMAT2 or SRSF1 was silenced using siRNAs and cells were grown for 5 days after which RNA was isolated and sequencing libraries prepared. Analysis was performed using DEXSeq. (B) Percentage of total detected differentially spliced, based on identified genes (left) or exonic regions (right) as determined with the DEXSeq package. (C) Differential exon usage, plotting the fold change over siControl vs siSRSF1 or siZMAT2 (N=3). (D) Differentially spliced genes after knockdown of ZMAT2 are enriched for binding of ING5 (ChIP-seq data (Mulder *et al.*, 2012)) and keratinocyte master regulator TP63. GO term analysis on these genes shows enrichment for focal adhesion related terms.

### Functionally interacting genes are jointly regulating an expression program involved in cell adhesion

The above presented results, together with the network prediction and functional interaction experiments (Figure 2), suggest that there is a link between epigenetic regulation and ZMAT2-mediated splicing of genes involved in cell adhesion. Consistent with these functional interactions, we found that the transcripts that are affected in their exon usage upon ZMAT2 silencing are highly enriched (p-value<10^−8^) in direct ING5 genomic binding targets which is one of the epigenetic factors in the functional interaction network (Figure 5D) (Mulder *et al.*, 2012) indicating that these genes regulate similar processes. In contrast, there was no evidence for regulation by DNA methylation or the heterochromatin associated H3K27me3 mark. Moreover, only a slight enrichment (p=0.003) for targets of the key epidermal transcription factor TP63 was found, suggesting that ING5 and ZMAT2 cooperate to regulate an RNA expression program important for epidermal biology. Notably, the gene-program targeted by both ZMAT2 and ING5 was further enriched in genes involved in focal adhesion formation, a process that is required to anchor epidermal stem cells to their niche. We conclude that joint epigenetic and splicing regulation of specific subsets of genes maintains epidermal stem cells in a proliferative, undifferentiated state.

## Discussion

Together, our results demonstrate that regulation of cell-cell adhesion related transcripts is at least orchestrated by a set of chromatin associated epigenetic regulators and a splicing component. We were able to discover their joint contribution by considering how genes cooperate in the regulation of gene expression programs, starting from our siRNA-based screens. Through these experiments we generated a rich dataset describing the roles of DNA/RNA binding factors and epigenetic regulators in human epidermal stem cell differentiation. The specific set-up of the experiments enabled us to use our previously published network Bayesian algorithm that predicts functional relationships between genes, transcending the annotation of a gene as a transcription factor, chromatin modifier or splicing factor, providing insight into how genes and processes work together to shape epidermal biology. Using this approach we uncovered a new functional connection between chromatin factors representing different epigenetic mechanisms and an uncharacterised zinc-finger, ZMAT2. Through proteomics and RNA sequencing approaches we validated that ZMAT2 is involved in RNA splicing in our system. More interestingly, we found that there is enrichment towards regulation of adhesion related genes through a RNA interaction experiments that suggests there is a targeted process involved. We did not find any evidence for specific deregulated splicing mechanisms that could explain this selectivity, nor could it be explained simply through transcript abundance.

It is conceivable that ZMAT2 functions through fine-tuning the splicing mechanism of a subset of genes. We reason that this is a likely scenario when considering the different steps splicing entails and the potential role for ZMAT2 in these. Splicing is an important post-transcriptional mechanism which catalyses the excision of non-coding parts in a pre-mRNA transcripts, or in *alternative* splicing the exon composition in a mature mRNA transcript. It is regulated by the very large and dynamic spliceosome complex, which facilitates the different steps of the splicing reaction through rearrangements in protein composition for each step. In the first step, the so-called complex A recognizes the splice site, after which complex B is assembled. This complex is catalytically activated to the B* complex, via the intermediate B^act^ complex. The B* complex catalyses the first transesterification reaction and where after more rearrangements generate complex C which catalyses the second transesterification reaction. The spliceosome is then disassembled and the subunits are recycled for a next round. ZMAT2 is an interactor of the B complex, were it accompanies the core complex B component tri-snRNP U4/U6.U5. In yeast, this tri-snRNP is already associated with ZMAT2 homolog snu23 and pre-mRNA processing factor 38 (Prp38) strictly prior to integration into the yeast pre-spliceosome. In contrast, in humans these two proteins are recruited independently from the tri-snRNP and are therefore considered non-snRNP proteins. There are seven more of these non-snRNP proteins in humans (Ulrich and Wahl, 2017) which are referred to as B-specific proteins. Although some of these proteins can influence splice-site selection (e.g. Smu1 and RED (Spartz, Herman and Shaw, 2004)) their exact function is not known. It is interesting to consider these proteins as mediators of alternative splicing through recognition of different splice-sites, which might explain the observed enrichment for adhesion related genes for ZMAT2. However, even though ZMAT2’s interaction with the human spliceosome is less stringently required than in yeast, it might have a more important function in enabling more general conformational changes from B to C complex in the spliceosome. In yeast, snu23 is thought to influence activation of the Brr2 helicase via its stabilisation on the U4 snRNA, which is part of the U4/U6.U5 tri-snRNP. The Brr2 mediated unwinding of the U4/U6 duplex is an important step towards generating a catalytically active spliceosome, because it frees the U6 snRNA to form an internal loop an helix with the U2 snRNA. This enables the conformational changes needed for the branching step and consequent exon ligation (Plaschka, Lin and Nagai, 2017). It is possible that ZMAT2 transiently interacts with the human spliceosome to facilitate and fine-tune these transitions. If this is the case, the timing of its association with the spliceosome might influence the efficiencies of competing splicing mechanisms, spliceosome composition or conformation (Ulrich and Wahl, 2017). Although we can only speculate about the molecular mechanisms behind our observations at this point, the combined evidence from our proteomic data and literature identified ZMAT2 as regulator of (alternative) splicing in epidermal stem cells.

Our results suggest that cell-cell adhesion is regulated by a previously unanticipated connection between transcriptional processes and post-transcriptional regulation. Experiments investigating the functional interactions between epigenetic modifiers and ZMAT2 revealed mostly alleviating interactions, stressing the importance of proper splicing in parallel with epigenetic control of adhesion related transcripts. We confirmed these connections experimentally by showing that the genes that were differentially spliced upon silencing of ZMAT2 were enriched for binding of ING5. Moreover, these genes were also enriched for binding of p63, indicating that the functionally connected genes are in charge of regulating a gene set that is essential for epidermal biology. Considering the importance of proper regulation of adhesion signalling for maintaining the stem cell fate (Levy *et al.*, 2000), it is not surprising that incorrect regulation leads to the severe effects observed upon silencing these genes individually. This in combination with the profound effect of silencing ZMAT2 individually on the differentiation state of the cells shows that the fine-tuning of splicing by ZMAT2 is of utmost importance for keratinocyte biology.

We have discovered that this group of epigenetic modifiers and ZMAT2 represent important hubs in the regulation of adhesion related transcripts and might be interesting to study more mechanistically in light of epidermal differentiation. We conclude that epigenetic and splicing factors jointly regulate processes that are essential for epidermal biology. Taken together, our work provides new insight into the regulation of epidermal differentiation and highlights the importance of cooperation among disparate cellular processes in the regulation of cell behaviour.

## Author contributions

S.T. designed, performed and analysed all experiments, P. J. and M.V. performed proteomics experiments K.M. analysed and interpreted data, conceived and oversaw the study.

## Acknowledgements

The authors would like to thank Dr. W. Mechelenbrink for advice and help on computational analyses and members of the Mulder lab for fruitful discussion. This work was supported by a VIDI grant from the Netherlands Organisation for Scientific Research (NWO-VIDI) to K.W.M.

---

**Supplemental figure 1:**
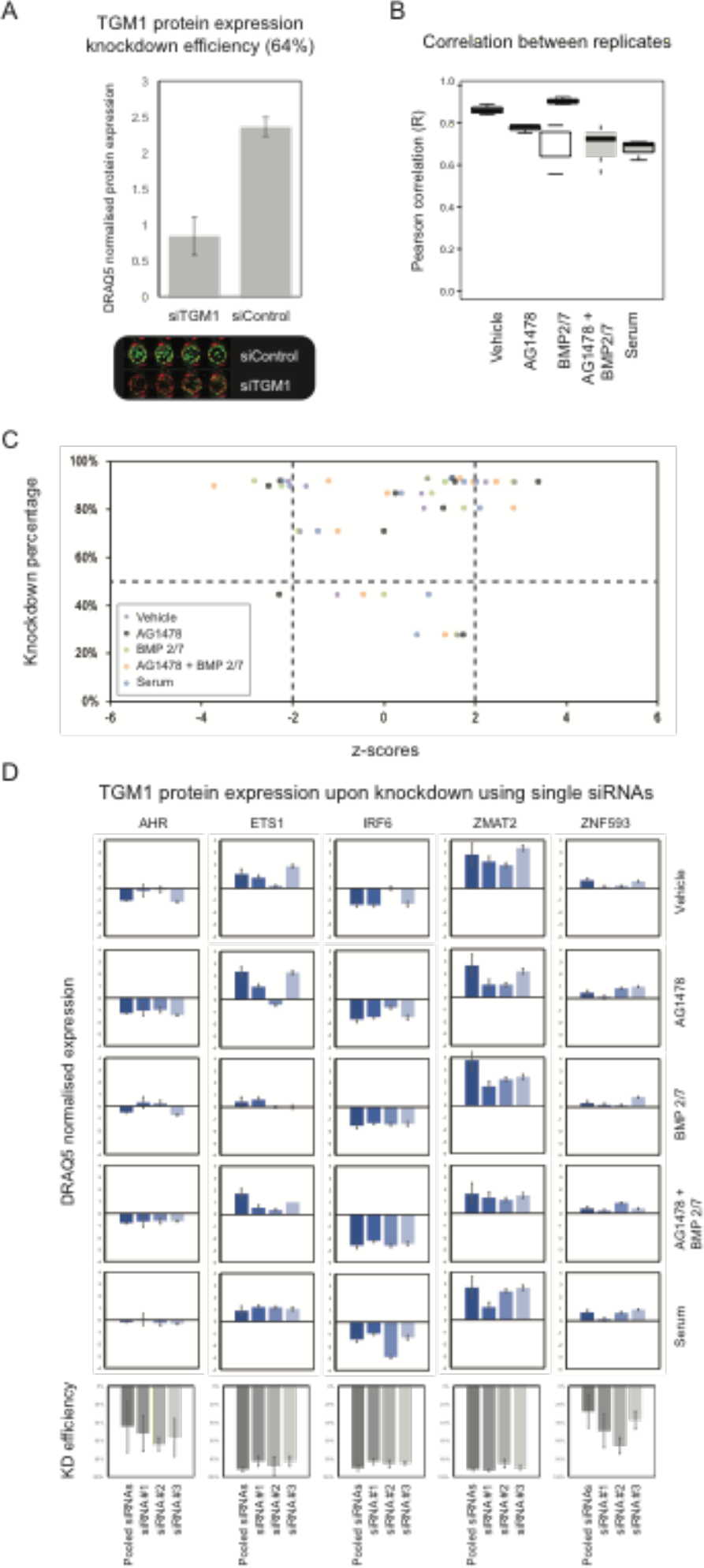
siRNA screen quality controls. (A) DRAQ5 normalised protein expression transglutaminase I after silencing using siRNAs (N=4, error bars indicate S.D.) showing antibody specificity. Knockdown efficiency: 64%. (B) Pearson correlation between replicate samples within one experiment for the different differentiation inducing conditions (N=3). (C) Estimation of the FDR, plotting the knockdown efficiency (percentage) versus the effects (z-scores) observed in the screen (N=3). (D) DRAQ5 normalised expression of TGM1 after silencing targeting the indicated genes in the 5 different conditions (N=3, error bars indicate S.D.). Knockdown efficiencies individual siRNAs vs pooled siRNAs targeting the indicated genes (N=3, error bars indicate S.D.).

**Supplemental figure 2:**
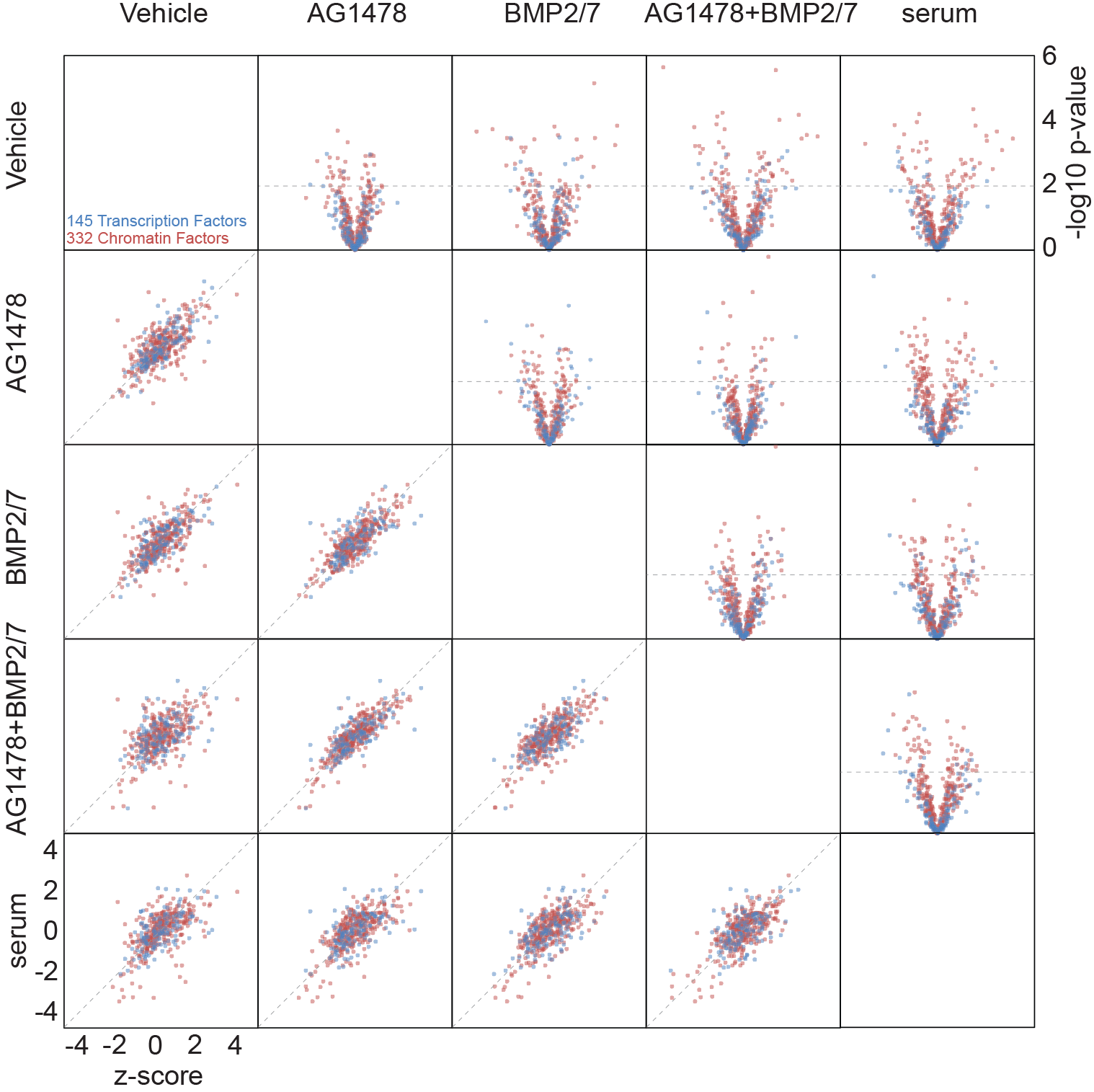
Conditional effects in the screen data suggest differential CF or NA binding factor dependencies during differentiation. Left-lower panels display correlation between replicate samples between conditions, and right-upper panels show effect size and significant differences between indicated conditions (N=3).

**Supplemental figure 3:**
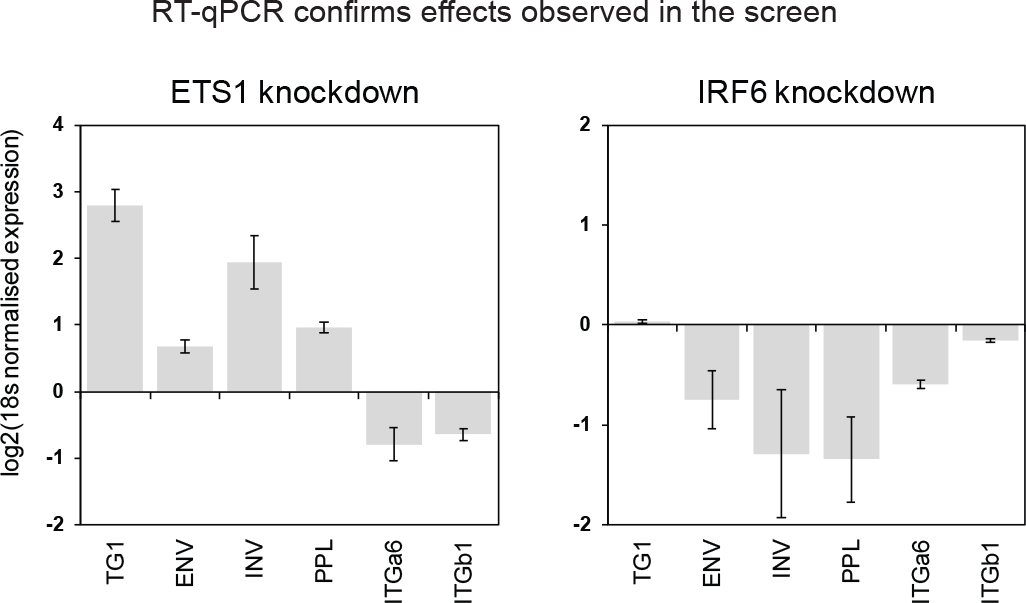
Functional validation of effects of silencing ETS1 and IRF6 on the differentiation state of the cell. (A) qPCR validation of expression of differentiation markers transglutaminase I (TGM1), envoplakin (ENV), involucrin (INV), periplakin (PLL) and self-renewal markers integrin alpha-6 (ITGa6) and integrin beta-1 (ITGb1) (N=3, error bars indicate S.D.). ZMAT2 KD percentage (72h): 91%, IRF6 knockdown percentage (72h): 92%.

**Supplemental figure 4:**
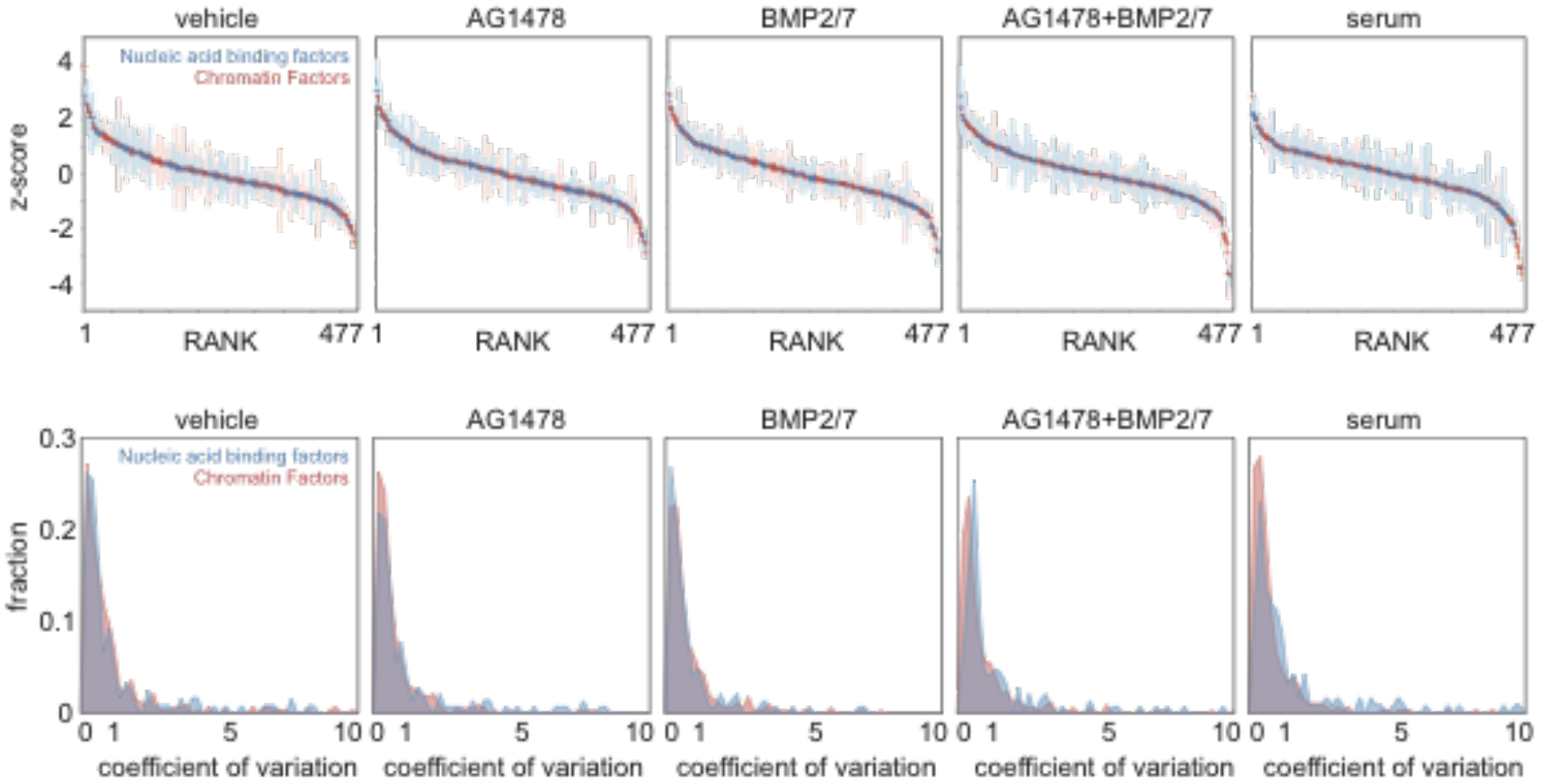
Data distribution of dataset describing the role of 332 chromatin factors and 145 NA binding factors. (A) Ranked z-scores and standard deviations for the combined dataset, with data origins coloured with NA binding factors in blue and chromatin factors in red (N=3, error bars indicate S.D.). (B) Histograms describing the distribution of variation of each dataset, with coefficient of variation of the chromatin factor dataset in red and the coefficient of variation of the NA binding factor dataset in blue (N=3).

**Supplemental figure 5:**
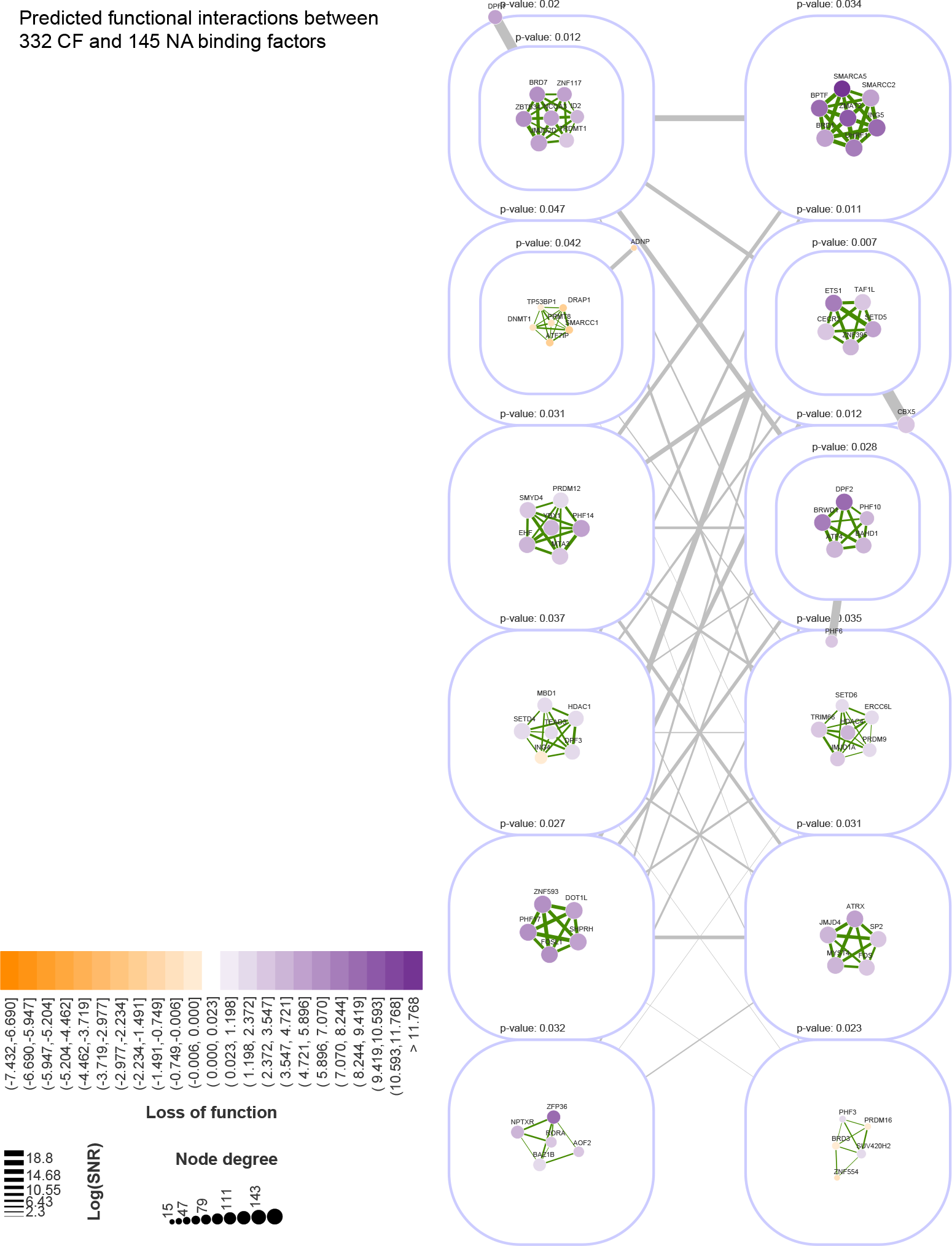
The complete functional interaction network as determined using the Bayesian mixture model. Purple lines circle enriched functional interactions and in edges in green indicate the SNR. Displayed significance indicates the approximately unbiased (AU) p-value after 10.000 rounds of bootstrapping.

**Supplemental figure 6:**
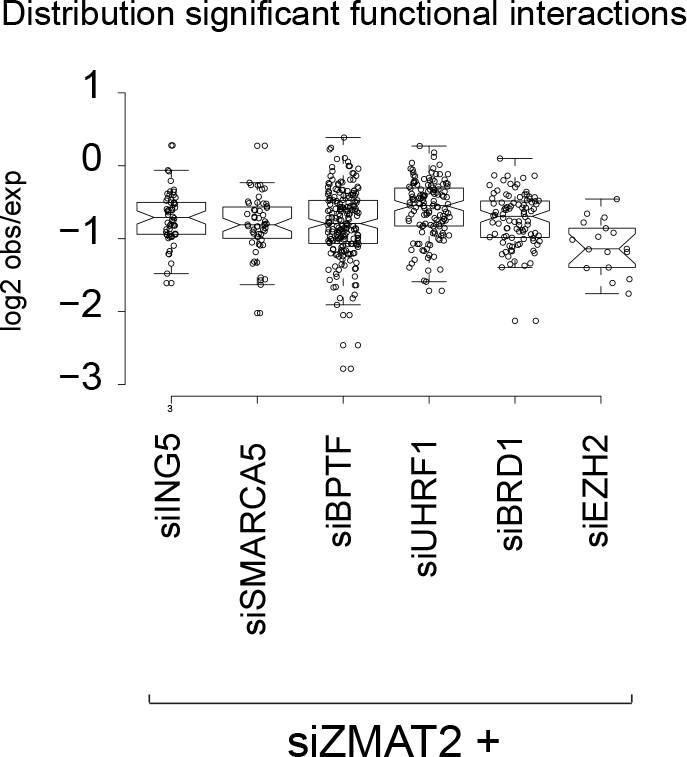
Data distribution functional interactions between ZMAT2 and epigenetic factors. Plot displays the logarithmic conversion of the ratio observed over expected effects for genes of which the observed expression is significantly different than expected for the indicated combinations. For siZMAT2 in combination with ING5 there are 57 genes differentially expressed than expected, for UHRF1 85, for BRD1 95, for SMARCA5 55, for BPTF 197 and for EZH2 17.

**Supplemental figure 7:**
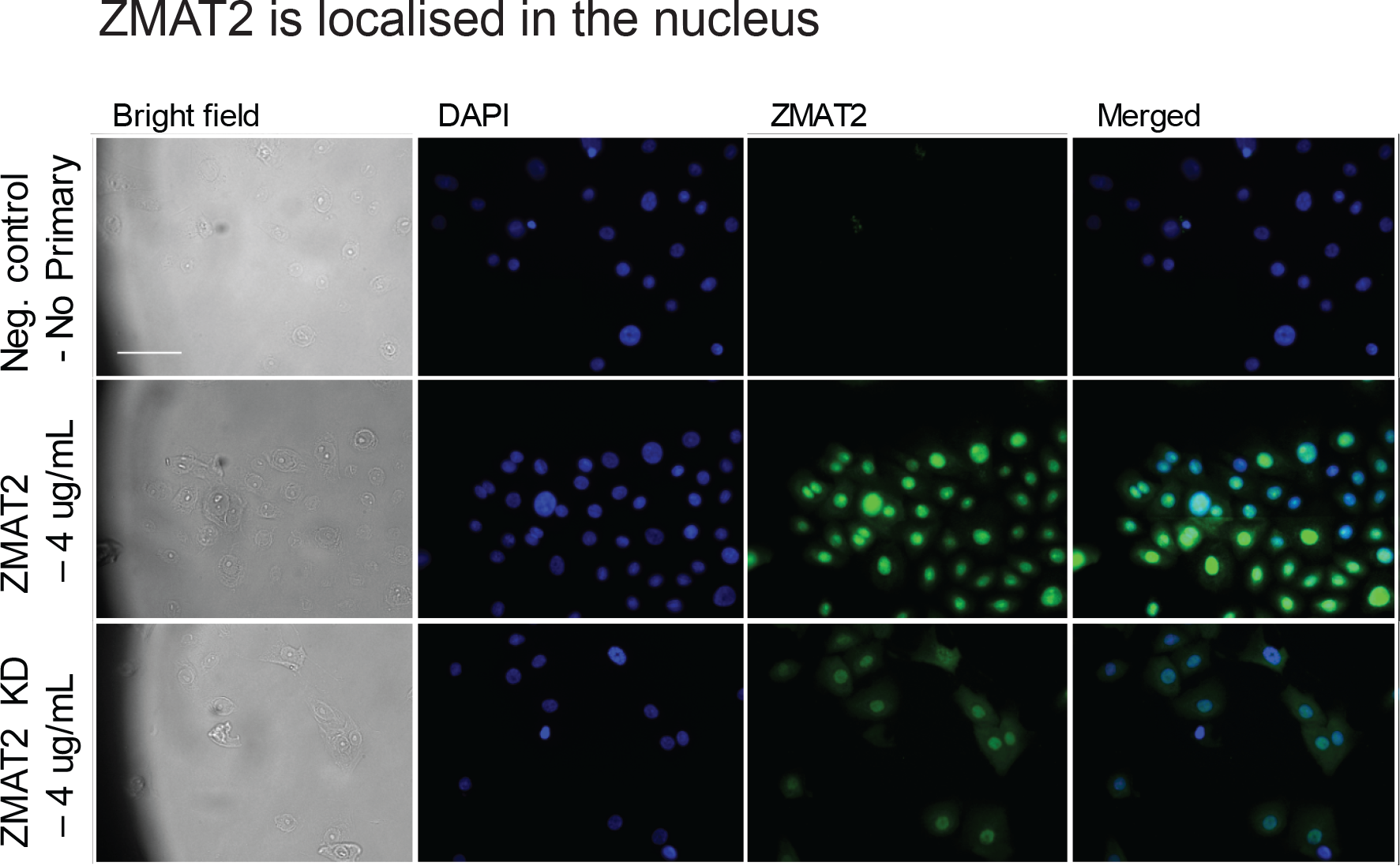
ZMAT2 is localised in the nucleus of human epidermal stem cells. Cells treated with mock siRNAs or ZMAT2 targeting siRNAs were stained with 4 μg/mL antibody supernatant and a no primary control sample was also included.

**Supplemental figure 8:**
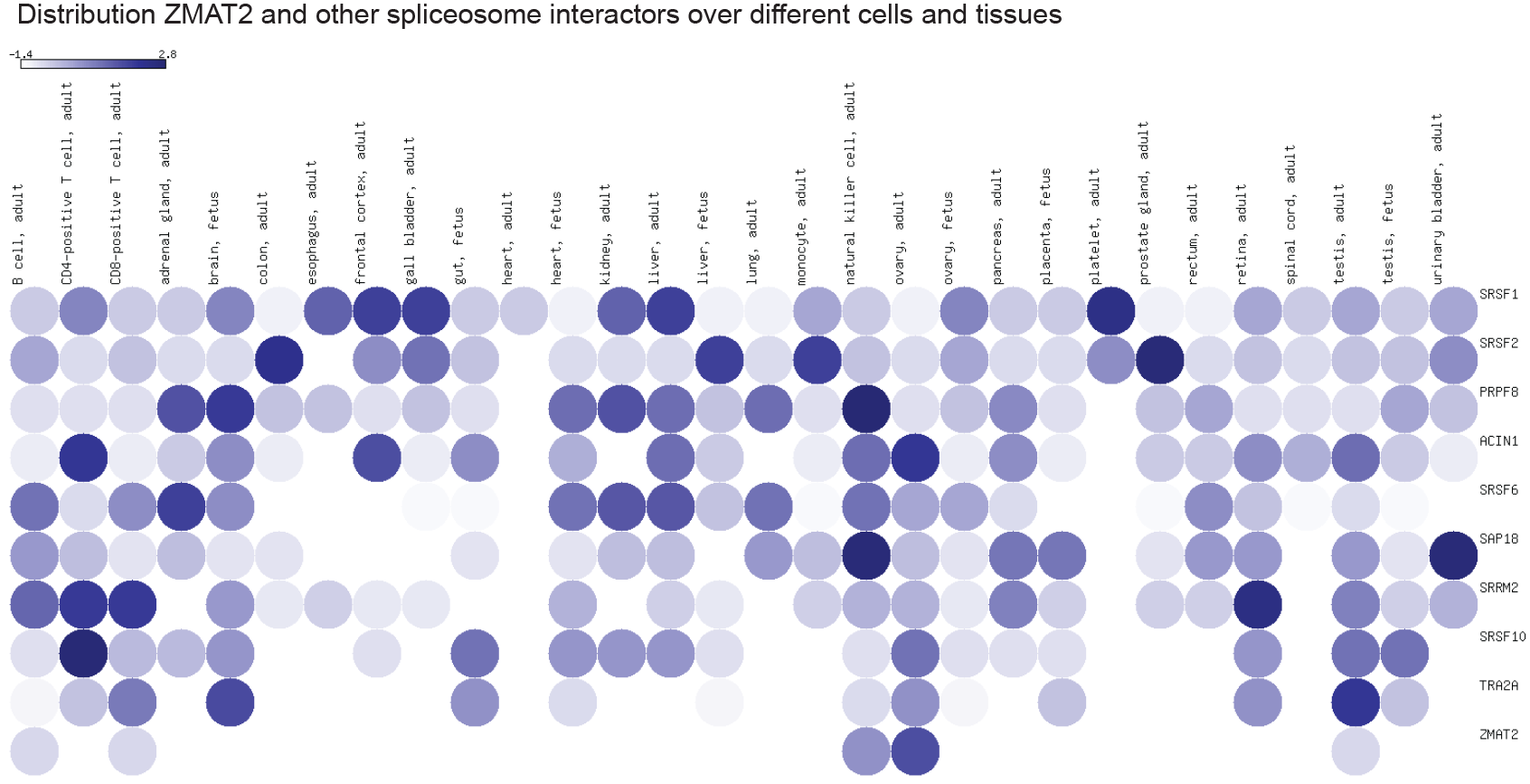
Distribution of expression of ZMAT2 and other spliceosome components over different cells and tissues in the human body. Colour of the circles indicates expression level if detected, as downloaded from the EMBL expression atlas.

**Supplemental figure 9:**
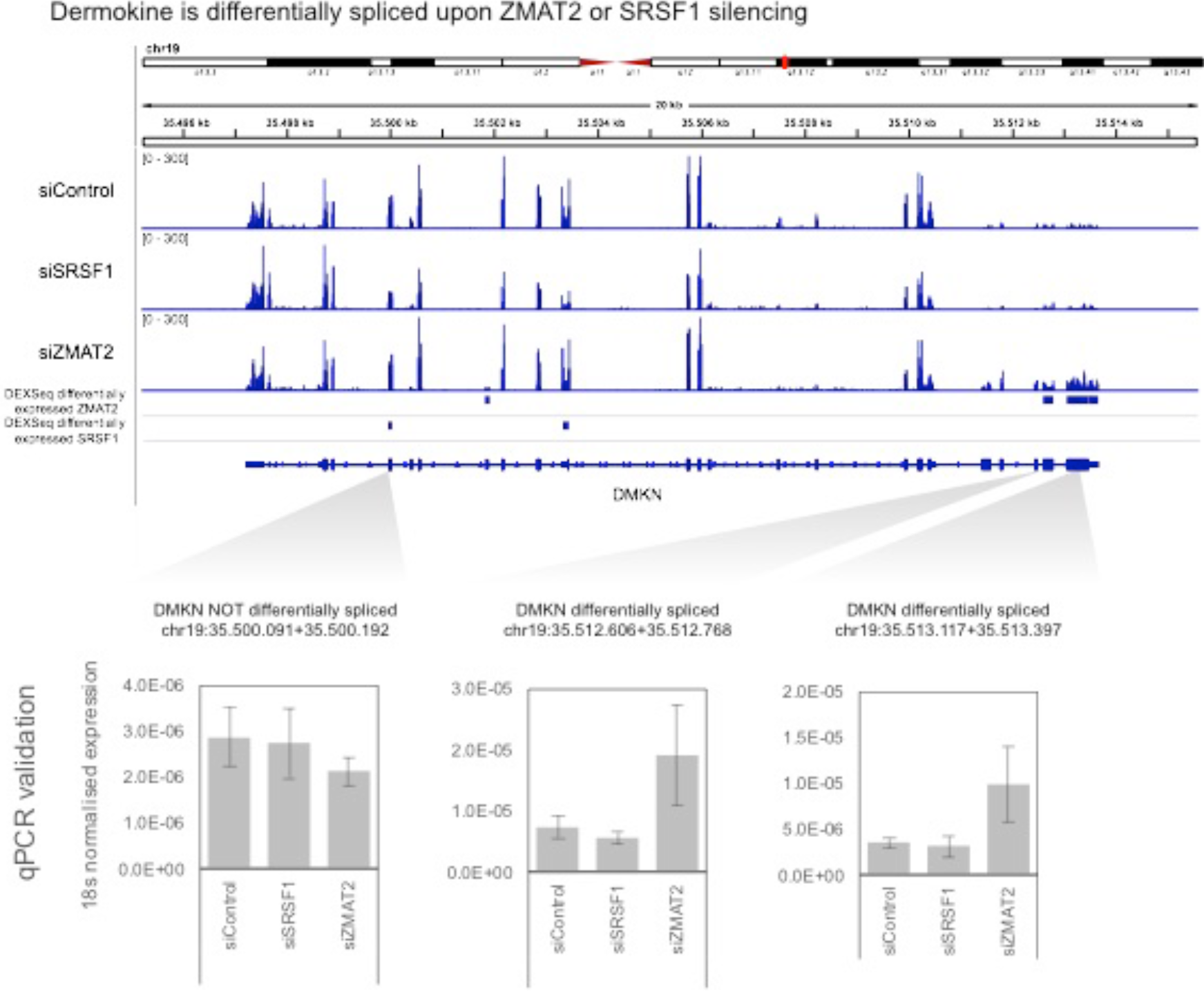
Differential splicing of DMKN upon silencing SRSF1 orZMAT2. (A) RNA sequencing tracks showing expression of DMKN exons upon silencing SRSF1 or ZMAT2, with below differential exons as called by DEXSeq. (B) qPCR validation of differential expression of exons (N=3, error bars indicate S.D.).

**Supplemental figure 10:**
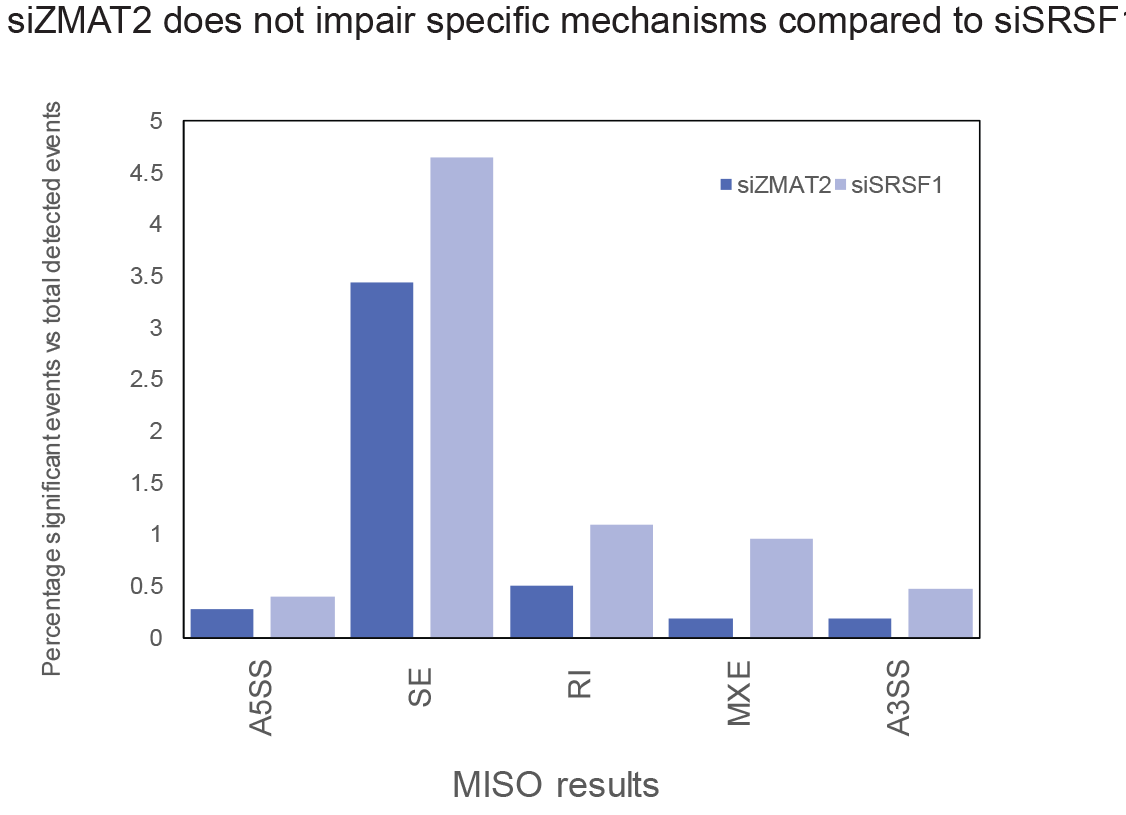
siZMAT2 does not impair specific mechanism compared to siSRSF1. Plot displays percentage significant events versus total detected events using the annotations provided on the MISO webpage under: Human genome (hg19) alternative events v1.0, describing skipped exons (SE), alternative 3’/5’ splice sites (A3SS, A5SS), mutually exclusive exons (MXE) and retained introns (RI).

**Supplemental figure 11:**
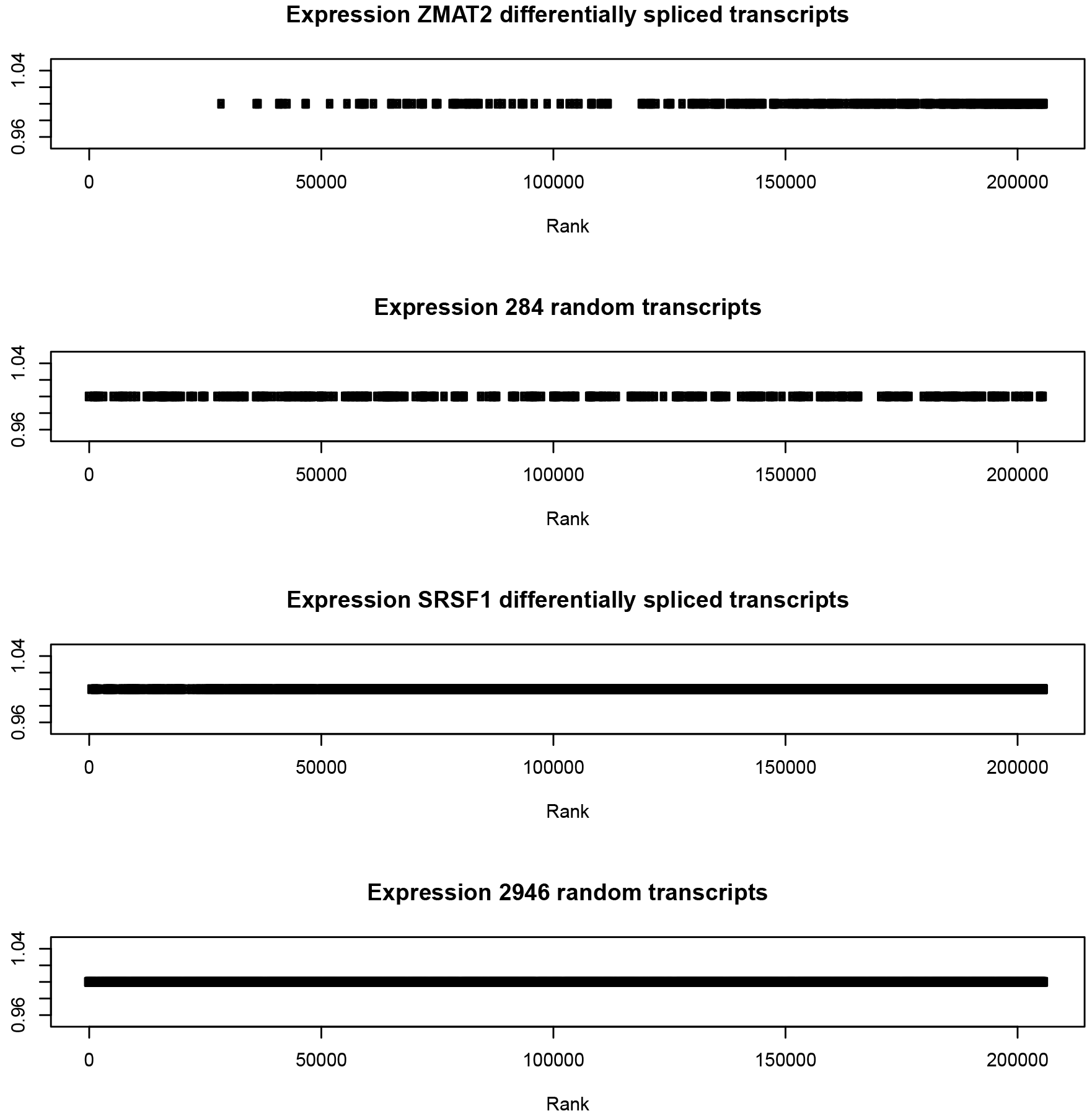
The abundance of genes that are differentially spliced upon silencing ZMAT2 or SRSF1. Plots display the ranked abundance of the 284 or 2946 genes that are differentially spliced upon knockdown of ZMAT2 or SRSF1. This is compared with 284 and 2946 randomly selected genes, respectively.

